# Dynamic assessment of the allocation of copper to cytochrome c oxidase using size-exclusion chromatography (SEC) combined with inductively coupled plasma mass spectrometry (ICP-MS)

**DOI:** 10.1101/2025.06.11.659066

**Authors:** Dina Secic, Megan E. Bischoff, Lucas Schmidt, Katherine E. Vest, John T. Cunningham, Julio A. Landero, Maria F. Czyzyk-Krzeska

## Abstract

Copper (Cu) is an essential trace element required for mitochondrial respiration via its incorporation into cytochrome c oxidase (CuCOX), the terminal enzyme of the electron transport chain. In this study, we employed size-exclusion chromatography coupled with inductively coupled plasma mass spectrometry (SEC-ICP-MS), UV-Vis spectroscopy, and immunoblotting to identify and validate a high-molecular-weight Cu-containing peak in SEC-ICP-MS chromatogram as representative of CuCOX activity. We demonstrate that this CuCOX peak is enhanced under metabolic conditions favoring oxidative phosphorylation, such as high Cu supplementation or galactose-containing media, and correlates with increased mitochondrial respiration. By tracing exogenously supplied ^63^Cu, we characterized the time- and dose-dependent incorporation of newly acquired Cu into CuCOX. Functional RNA interference (RNAi) experiments targeting key Cu transporters revealed that CuCOX formation is independent of the high-affinity Cu importer CTR1, but instead relies on alternative transporters including DMT1, LAT1, and the mitochondrial carrier SLC25A3. These findings offer new insight into the cellular pathways governing Cu trafficking and allocation to mitochondria under physiologically relevant conditions. Furthermore, our work establishes SEC-ICP-MS as a sensitive and specific method for quantifying CuCOX and assessing mitochondrial metabolism. This platform holds promise for the identification of Cu-related biomarkers and therapeutic targets, particularly in the context of diseases such as renal cell carcinoma (RCC), where dysregulated Cu homeostasis plays a critical role.

## Introduction

Copper (Cu) is an essential trace element involved in a wide range of biological processes, including mitochondrial energy production and the function of various enzymes. One of copper’s most vital roles is in cytochrome c oxidase (CuCOX), the terminal enzyme and rate-limiting step of the mitochondrial electron transport chain. CuCOX facilitates the transfer of electrons to molecular oxygen, a crucial process that drives ATP production. The enzyme contains two essential copper centers: CuA and CuB. The CuA site, a binuclear copper center, transfers electrons from cytochrome c to the enzyme’s catalytic core, while the CuB site, together with a heme a3 group, forms a dinuclear center that binds molecular oxygen and catalyzes its reduction to water (1, 2). The assembly and activity of the CuCOX complex are tightly regulated by Cu availability.

The intracellular uptake, distribution, and speciation of copper are tightly regulated to ensure proper cellular function and prevent toxicity (3). Extracellular Cu is imported through a plasma membrane high-affinity transporter, CTR1, which is internalized into endosomes in response to high Cu to protect cells from Cu toxicity (4). CTR1 transport requires reduction of the extracellular Cu^2+^ to Cu^+^ by STEAP metalloreductases. In some cells, Cu^2+^ is imported by a divalent metal transporter, DMT1 (SLC11A2) (5) or bound to histidine and imported by the amino acid transporter SLC7A5 (LAT1) (6). One study reported macropinocytosis as an exclusive Cu uptake mechanism in RAS-driven colon cancer tumors (7). Intracellular Cu is bound to metallothioneins (MTs) for storage (8) and to protein chaperones for transport to cuproenzymes (9). The labile, dynamic pool of Cu is bound to small molecules (e.g. GSH) and contributes to the allosteric effects of Cu. Cu enters mitochondria through the SLC25A3 transporter (10, 11) and is delivered for the formation of CuCOX by COX17 to downstream chaperones COX11/COX19 (2) and SCO1 that respectively transfer Cu to the CuB center in the COX subunit 1 and the CuA center in the COX subunit 2 (1). ATP7A/B transporters function as the primary Cu detoxification pathway in eukaryotic cells (12), and export Cu to Golgi for assembly into secreted cuproenzymes and into the extracellular environment at the plasma membrane (3).

Our recent studies showed that elevated levels of Cu and CuCOX are hallmarks of tumor progression in clear cell renal cell carcinoma (ccRCC) (13). Canonically, ccRCC has high glycolytic and low mitochondrial activity due to the orchestrated transcription program resulting from activation of hypoxia inducible factors (HIFs) due to the loss of the von Hippel-Lindau tumor suppressor (VHL) (14). However, more aggressive or relapsing tumors accumulate Cu and allocate it to CuCOX, leading to a major shift in metabolic state towards oxidative phosphorylation and away from glycolysis and driving tumor growth (13, 15).

Size-exclusion chromatography (SEC) coupled with inductively coupled plasma mass spectrometry (ICP-MS) is a powerful technique for analyzing metal-protein interactions in biological samples (16). SEC enables the separation of biomolecules based on their size, and with the proper selection of the mobile phase preserves their native state, while ICP-MS provides highly sensitive and specific detection of metal content at each eluting fraction. A diode array UV-Vis detector can be added between SEC column and the ICP-MS detector as a non-destructive detector, providing information related to chromophores typical of metal-binding proteins. This combination allows for a molecular mass-based characterization of metalloproteins, their metal-binding properties, and potential changes under physiological or pathological conditions.

Using orthogonal approaches, we established that the high-molecular-weight peak detected in Cu SEC-ICP-MS chromatograms represents an active fraction of CuCOX and can be used to monitor the dynamic interplay between Cu availability, metabolic states, and CuCOX activity. Notably, we found that the incorporation over 48 hoursof exogenous ^63^Cu is required for CuCOX induction in renal cancer cells. This process is independent of the high-affinity Cu importer CTR1 and instead relies on DMT1, the amino acid transporter LAT1, and the mitochondrial carrier SLC25A3. Our findings position ICP-MS and SEC-ICP-MS as powerful tools for studying Cu uptake and CuCOX biogenesis and activity, with potential applications in analyzing primary tumors and patient sera to explore Cu and CuCOX as prognostic biomarkers in clear cell renal cell carcinoma.

## Results

### High Molecular Weight Peak in SEC-ICP-MS chromatogram represents CuCOX activity

Total non-denatured lysates from renal carcinoma cells exposed for 48 h to standard media containing 0.14 µM Cu or media supplemented with 30 µM Cu were analyzed by SEC-ICP-MS. Peaks were identified based on their retention times, signal intensity, and comparison with mass calibration standards.

A standard Cu SEC-ICP-MS chromatogram includes the heavy molecular weight fraction (HMW, 5-8 min retention time) that measures Cu bound to proteins; Cu bound to the cysteine-rich storage proteins, metallothioneins (MT, 8-10 min retention time); and the low molecular weight fraction (LMW, after 12 min) of Cu bound to metabolites, such as GSH, which reflects labile Cu (Fig. 1*A*, *top*). Importantly, a peak with retention time at 5-6 min represents mitochondrial CuCOX:

**Figure 1.**
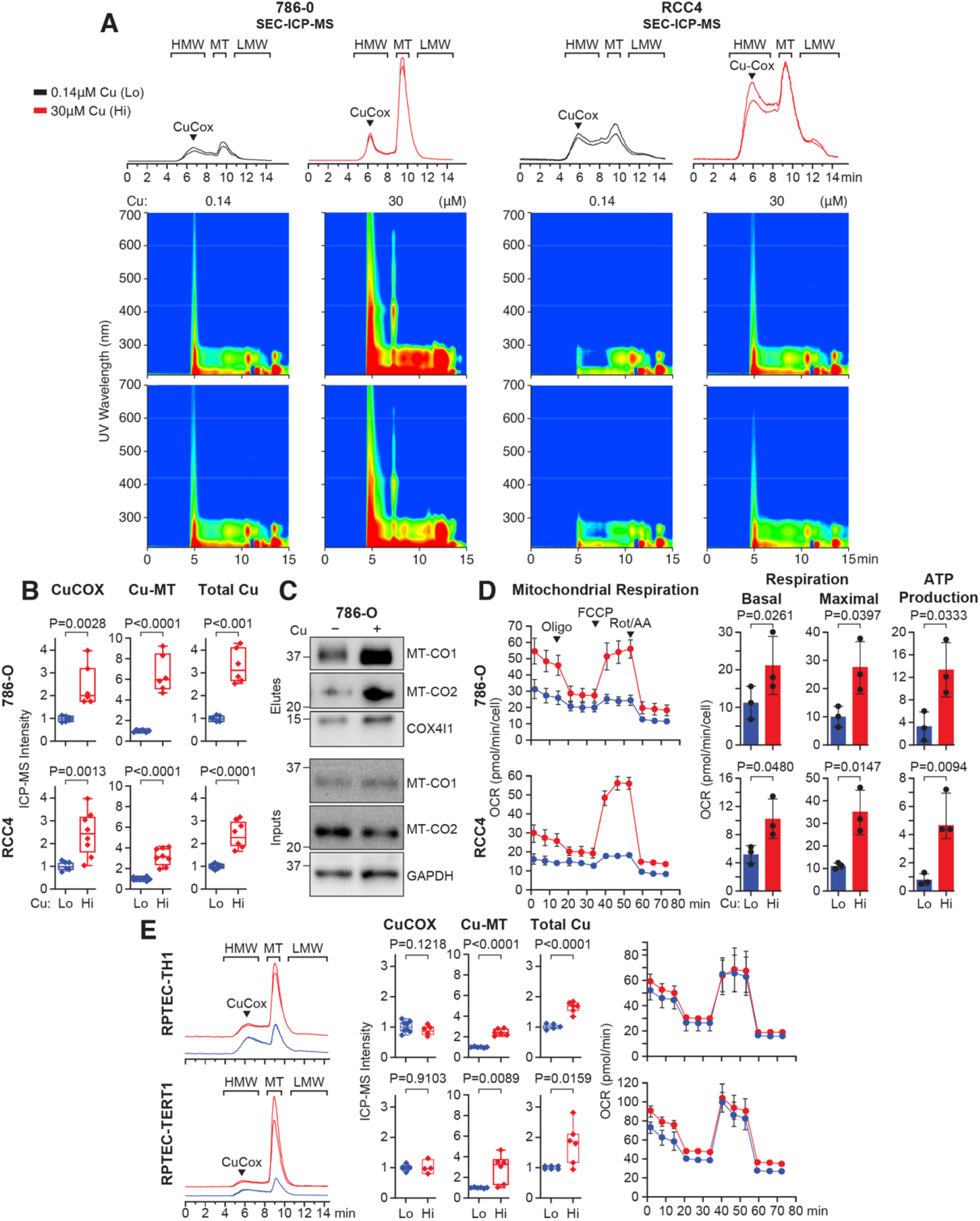
Identification and validation of CuCOX peak in SEC-ICP-MS chromatogram. *A*, Top: Representative SEC-ICP-MS chromatograms of lysates from 786-O and RCC4 cells grown in 0.14 µM or 30 µM Cu for 48 h. Bottom: UV-Vis absorbance isoplots collected throughout the entire chromatography run. Note that, because absorbance was measured before the ICP-MS detector, there is a slight shift in the timeline between the chromatogram and the spectra. HMW: high molecular weight fraction, MT: metallothioneins, LMW: low molecular weight fraction. *B*, Box whisker plots show relative abundance of Cu measured as ICP-MS intensity for Cu allocated to CuCOX, MTs and total Cu content in 786-O (3 biological and 2 technical replicates) and RCC4 (4 biological and 2 technical replicates) cells. *C*, Western blot of indicated proteins eluted from the CuCOX peak and the inputs are shown below. *D*, Oxygen consumption rate (OCR) from representative seahorse mitochondrial stress tests in 786-O and RCC4 cells (3 biological replicates). Quantification of basal and maximal respiration (post-FCCP injection), and respiration coupled to ATP production. Means±SD are shown. P value for basal respiration in 786-O cells was calculated using a paired two-tailed t-test. *E*, SEC-ICP-MS chromatograms, quantification of CuCOX and MT peaks, total Cu level, and Seahorse measurements of OCR for RPTEC TH1 and TERT1 cell lines (3 biological replicates and 2 technical replicates for each). Box-and-whisker plots display median, minimum and maximum values and all individual data points. Unless otherwise indicated, P values were determined using a two-tailed t-test.

This peak is induced by treatment of 786-O and RCC4 renal cancer cells with media containing 30 µM Cu for 48 h (Fig. 1, *A* and *B*). Analysis of the UV-Vis absorbance isoplots in parallel to the SEC-ICP-MS reveals increased absorbance of the spectrum in response to Cu exposure at the 6 min time point corresponding to CuCOX peak in the chromatogram (Fig. 1*A*, bottom). This includes 420-450 and 600 nm absorbance bands corresponding to cytochrome c oxidase due to the presence of heme a and a3 (17-20) (Fig. 1*A*, bottom). The absorbance peak detected at 7 min is not detected by SEC-ICP-MS as it does not contain Cu. Western blot analysis of eluates from the fraction corresponding to CuCOX peak shows Cu-induced enrichment for mitochondrially encoded, MTCO1 and MTCO2 subunits in response to Cu treatment (Fig. 1*C*). Exposure to 30 µM Cu also increased Cu bound to MTs and the total Cu content in both cells (Fig. 1*B*). Functionally, exposure to 30 µM Cu resulted in enhanced mitochondrial oxygen consumption rate (OCR) and ATP production in 786-O and RCC4 RCC cells that have constitutively low OCR (Fig. 1*D*). Furthermore, to compare the effects of Cu in renal cancer cells with non-cancer cells, we expanded this analysis to two immortalized Renal Proximal Tubule Epithelial Cell (RPTEC) TH1 and TERT1. Interestingly, both RPTEC cell lines accumulated higher levels of Cu bound to MTs, but there was no effect on the CuCOX peak or OCR, possibly because these cells had higher basal respiration as compared to RCC cells (Fig. 1, *E*). These data indicate that Cu regulates mitochondrial electron transport chain and CuCOX differently in renal cancer vs. epithelial cells at least during 48-hour exposure.

Next, we investigated whether oxidative phosphorylation-inducing conditions, beyond Cu overload, influence CuCOX dynamics as detected by SEC-ICP-MS. In that respect, replacing glucose with galactose in the tissue culture media forces cells to rely on oxidative phosphorylation (21, 22). However, the biochemical mechanism of this established experimental observation is not clear. One possibility is that the conversion of galactose to glucose through the Leloir pathway slows glycolytic generation of ATP and therefore requires activation of oxidative phosphorylation (Fig. 2*A*). Indeed, 48 h exposure of 786-O cells to media that contains 0.14 µM Cu and 10 mM galactose significantly increased OCR and mitochondrial ATP production as compared to the same media with 10 mM glucose. However, the total production of ATP was smaller in cells grown in galactose as compared to glucose containing media (Fig. 2*B*). Consistently, with this functional readout, there was also an increase in CuCOX peak in SEC-ICP-MS chromatogram and corresponding UV-Vis absorbance isoplots (Fig. 2*C*). Surprisingly, not only was there an increase in Cu allocation to CuCOX, but there was also a significant induction of Cu bound to MTs and a total increase in Cu content (Fig. 2*D*), although these increases were smaller as compared to the exposure to 30 µM Cu as shown in Fig 1. This indicates that a demand for CuCOX biogenesis can regulate the entire landscape of Cu uptake and allocation. The eluate from the CuCOX peak in cells grown in galactose media showed enrichment of CuCOX subunits, MT-CO2, and COX4l1 (Fig. 2*E*). This galactose-induced increase in Cu was abolished by treating cells with inhibitors not only of CuCOX but also of electron transport chain complex I and III (Fig. 2, *F* and *G*), indicating coordinated regulation between overall ETC activity and Cu uptake and allocation.

**Figure 2.**
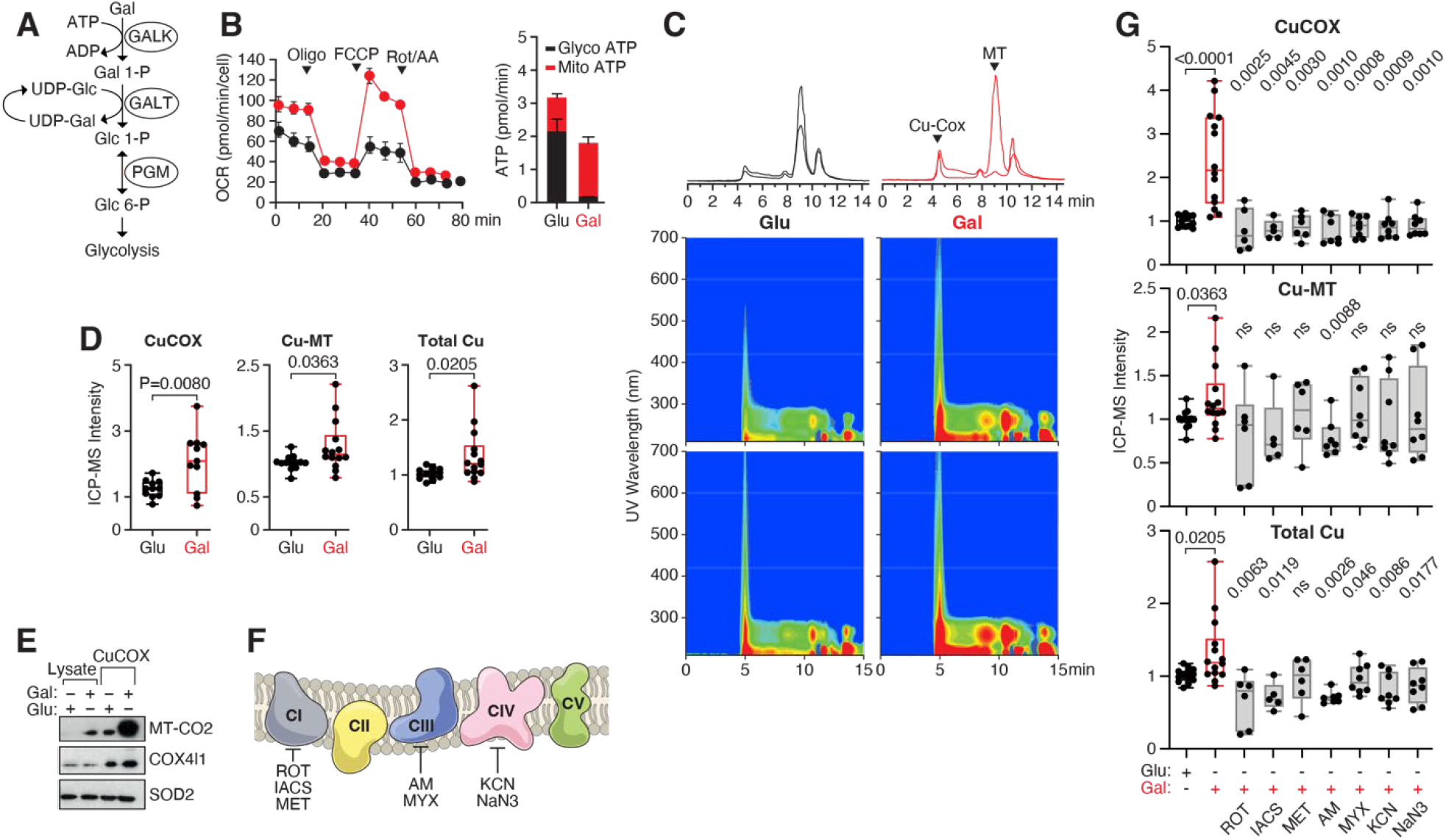
Galactose induces oxidative phosphorylation and increases Cu content and allocation to CuCOX in 786-O RCC cells. *A*, Schematic representation of the Leloir pathway converting galactose into glucose. *B*, OCR and mitochondrial ATP production in response to 48 h treatment with galactose (red) as compared to glucose (black) (three biological replicates). *C*, SEC-ICP-MS chromatograms of Cu (top) and corresponding UV-vis absorbance plots in lysates from 786-O cells treated with 10 mM glucose or galactose for 48 h. *D*, Box whiskers show quantification of Cu allocated to CuCOX and MTs, and total Cu content in lysates from 786-O cells grown in glucose- or galactose-containing media (7 biological replicates with two technical replicates for each). *E*, Western blot of total input lysates and CuCOX eluates from 786-O grown in glucose or galactose media. *F*, Schematic model of mitochondrial electron transport chain along with localization of inhibitors for each complex. *G*, Quantification of SEC-ICP-MS peaks corresponding to Cu in CuCOX and MTs, and total Cu in lysates of 786-O cells treated with media containing glucose, galactose, or galactose with the indicated electron transport chain inhibitors added for 3 h before collection. ROT, rotenone (200 nM, 3 biological replicates in technical duplicates), IACS, IACS-10759 (1nM, 3 biological replicates in technical duplicates), MET, metformin (25 mM, 3 biological replicates in technical duplicates), AM, antimycin A (1nM, 4 biological replicates in technical duplicates), MYX, myxothiazol (1 nM, 4 biological replicates run in technical duplicates), KCN, potassium cyanide (0.5 mM, 4 biological replicates in technical duplicates), NaN_3_, sodium azide (0.5 mM 4 biological replicates in technical duplicates). Box- and-whisker plots displayed the median, minimum and maximum values, and all individual data points. All P values were calculated using a two-tailed t-test.

### Kinetics of ^63^Cu incorporation into CuCOX complex

The natural relative abundance of the Cu stable isotopes, ^63^Cu and ^65^Cu, is 69.17% and 30.83%, respectively, reflecting an approximate 2:1 ratio maintained in naturally occurring Cu samples (25). This allows for the use of exogenous pure ^63^Cu to trace Cu uptake and allocation by monitoring both ^63^Cu and ^65^Cu and their ratio by SEC-ICP-MS, and at the same time, to monitor the fate of endogenous Cu (Fig. 3*A*). Here, we traced the incorporation of exogenous ^63^Cu into CuCOX over time. Cells were cultured in standard media containing 0.14 µM Cu (natural isotopic abundance) and then switched to media supplemented with 0.14, 10 or 30 µM ^63^Cu only for 2 to 48 h (Fig. 3*B*). 0.14 µM represents Cu concentration in DMEM/F12 media with 10% serum used for carrying the cells, while 10 µM Cu represents physiological Cu level in sera from healthy human subjects, and 30 µM Cu represents concentrations of Cu measured in sera from patients with ccRCC (13). ^63^Cu concentration had a major effect on the magnitude of ^63^CuCOX and MT peaks, with almost immediate allocation of Cu to MTs and slower to CuCOX (Fig. 3, *C-F*). There were also some differences between 786-O and RCC4 cells. The total uptake of Cu and its allocation to MT and CuCOX were significantly larger in 786-O cells and these cells were very sensitive to the increase in ^63^Cu between 10 and 30 µM (Fig. 3,*C* and *E*). In contrast, in the case of RCC4 cells, only the amount of ^63^Cu in the CuCOX peak was sensitive to the change in Cu concentration between 10 and 30 µM, while the total ^63^Cu uptake and ^63^Cu bound to MTs were the same (Fig. 3, *D* and *F*). RCC4 cells accumulated less ^63^Cu as compared to 786-O cells, which is consistent with our previously published findings (13). This can be related to the fact that 786-O cells are a malignant RCC cell line that expresses only HIF2A, while RCC4 cells are less malignant and express HIF2A and HIF1A, which is considered a tumor suppressor (26). Note that 48 h of exposure to high Cu is also the time point where we consistently measured induction of OCR (Fig. 1*D*). Shorter exposures to Cu did not result in a consistent increase in OCR (data not shown).

**Figure 3.**
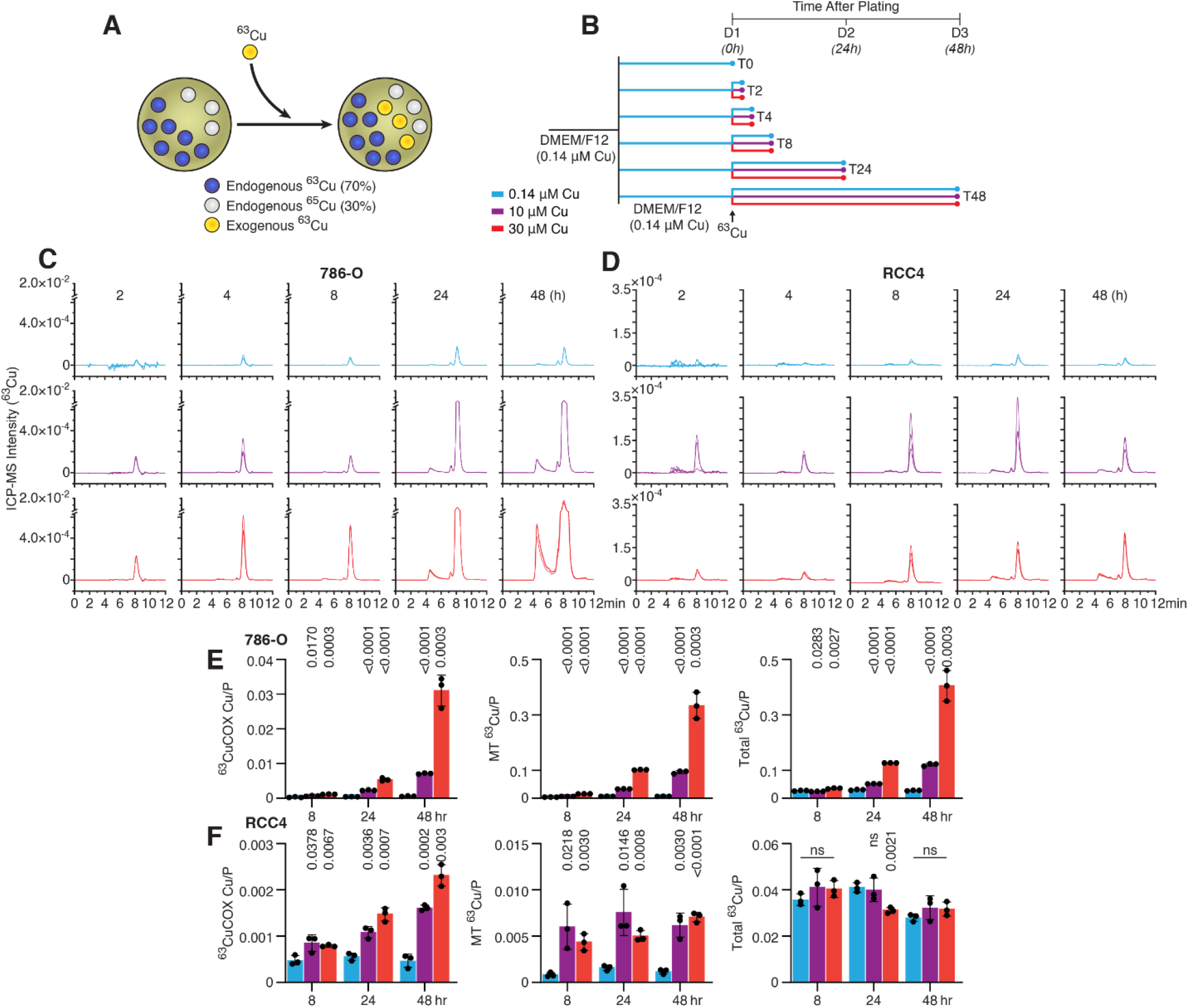
Kinetics of allocation of ^63^Cu tracer into CuCOX. *A*, Schematic representation of tracing ^63^Cu. *B*, Timeline of the tracking of ^63^Cu into CuCOX. *C* and *D*, SEC-ICP-MS chromatograms of exogenous ^63^Cu in lysates from 786-O and RCC4 cells at the indicated time points. *E* and *F*, Quantification of ^63^Cu in CuCOX, MTs and total exogenous ^63^Cu in 786-O and RCC4 cells. Cu abundance was normalized to Phosphorus (P). In all experiments: n=4, SEM±SD are shown; P values were calculated using a two-tailed t test and indicate significance of the difference between 10 or 30 µM Cu compared to 0.14 µM Cu. Experiments were performed in three technical replicates twice and a representative experiment is shown.

Next, we determined the effects of exposure to the exogenous ^63^Cu on the levels of intracellular exogenous ^63^Cu and endogenous Cu, Zn, and Fe. 786-O and RCC4 cells, which were chronically grown either in standard low Cu media (0.14 µM) or high Cu media (30 µM), were treated with 30 µM ^63^Cu for 48 h (Fig. 4*A*). Cells chronically adapted to high Cu concentration started with more endogenous Cu and accumulated more exogenous ^63^Cu as compared to cells chronically grown in low Cu media (Fig. 4B and C), which is also true for the Cu allocated to CuCOX (Fig. 4, *B* and *D*). Interestingly, in 786-O and RCC4 cells chronically grown in low Cu media, the uptake of exogenous ^63^Cu during 48 h of exposure to 30 µM Cu resulted in a decrease in endogenous CuCOX, and in the case of 786-O cells, also in total endogenous Cu (Fig. 4, *B* and *C, D*). This effect was not observed in cells grown chronically in high Cu media, indicating an early protective response, potentially to limit the amount of Cu during more acute exposures. In both cell lines, exposure to Cu affected levels of Zn (Fig. 4, *E* and *F*). Finally, cells chronically exposed to Cu showed increased Fe levels (Fig. 4, *G* and *H*). This is consistent with the role of iron in heme and iron-sulfur cluster proteins necessary for the ETC and CuCOX activities (27).

**Figure 4:**
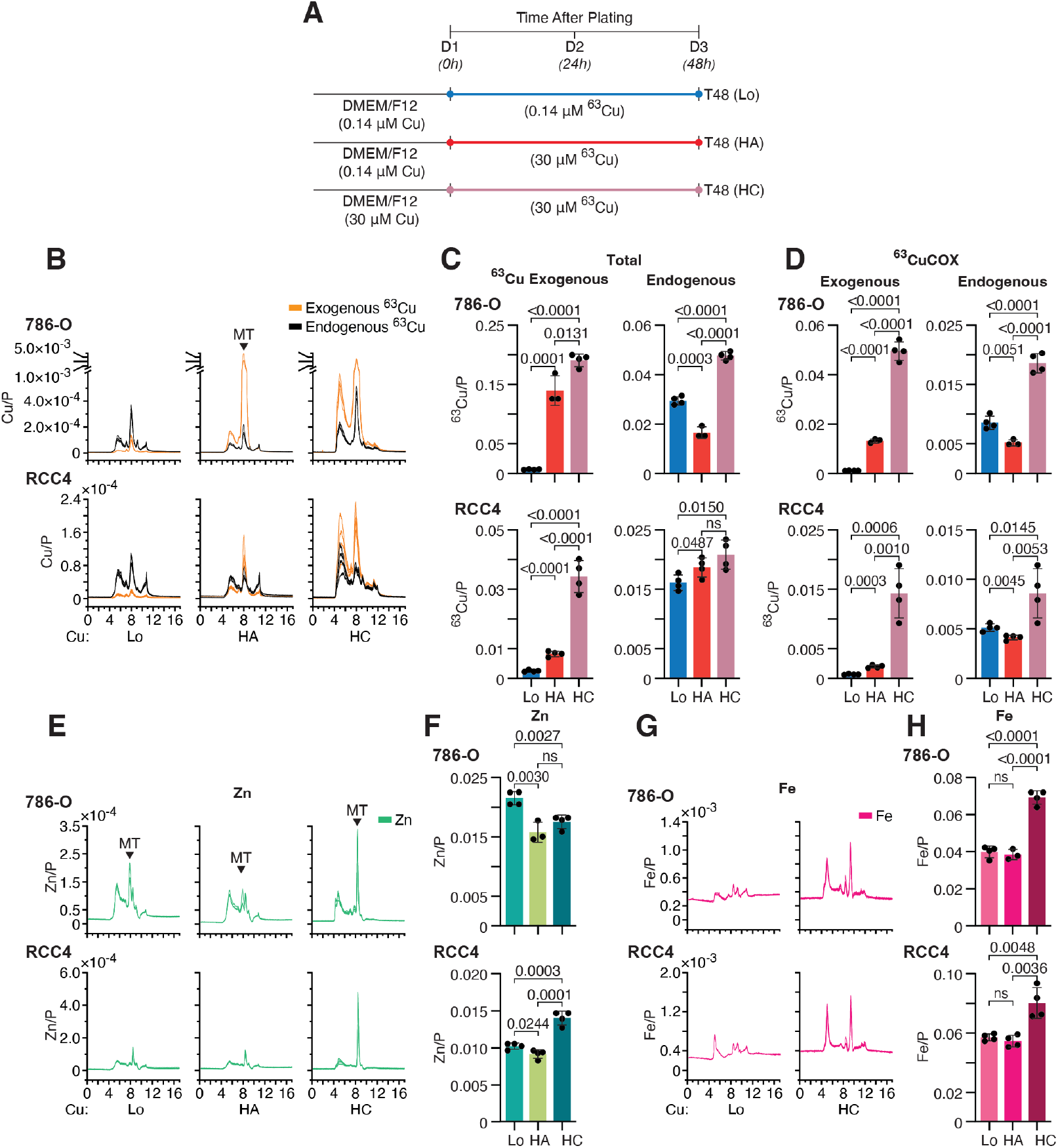
Allocation of tracer ^63^Cu to CuCOX in 786-O and RCC4 cells chronically grown in low or high Cu media. *A*, Timeline of the experiment: HA, exposure to 30 µM ^63^Cu for 48 h of cells grown continuously in low Cu media, HC, exposure to 30 µM ^63^Cu for 48 h of cells grown in high Cu media. *B*, Representative examples of Cu SEC-ICP-MS chromatograms from cells grown in Cu low media (Lo) and exposed to 30 µM ^63^Cu for 48 h (HA) or cells grown continuously in media containing 30 µM Cu and then treated with the same concentration of ^63^Cu. *C*, Quantification of total exogenous ^63^Cu and total endogenous Cu from SEC-ICP-MS chromatograms *D*, Quantification of ^63^Cu and endogenous Cu in CuCOX peak from SEC-ICP-MS chromatograms. *E*, Representative examples of Zn SEC-ICP-MS chromatograms as described in B. *F*, Quantification of total cellular Zn based on SEC-ICP-MS. *G*, Representative examples of Fe SEC-ICP-MS chromatograms as described in B. *H*, Quantification of total cellular Fe based on SEC-ICP-MS chromatograms. Metals’ abundance was normalized to Phosphorus (P). In all experiments: Means±SD are shown; n=4 biological replicates; P values were calculated using two-tailed t-test.

### Analysis of transporters contributing Cu for CuCOX

Extracellular Cu is primarily taken up by the high-affinity transporter, CTR1, but other transporters such as DMT1 and LAT1 have also been proposed (5, 6). Cu enters mitochondria via the phosphate transporter SLC25A3 (10, 11). To determine the role of these transporters in allocating exogenous ^63^Cu to CuCOX in cells grown in chronic low and high Cu conditions, we knocked down CTR1, DMT1, LAT1 and SLC25A3 with siRNA. Cells were collected 72 h after the first siRNA transfection and 48 h after exposure to exogenous 30 µM ^63^Cu (Fig. 5, *A* and *F*).

**Figure 5:**
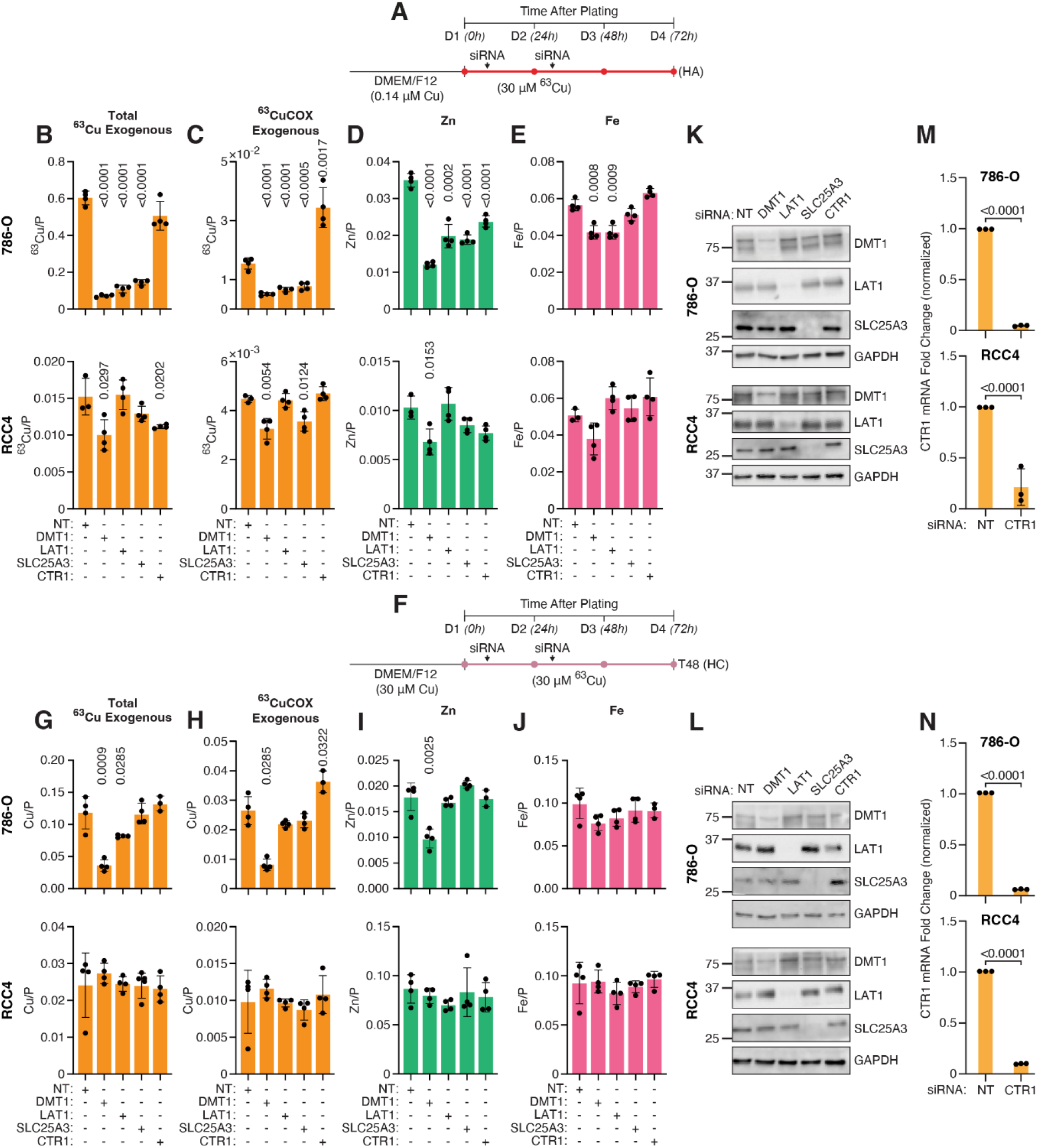
Contribution of Cu transporters to the allocation of tracer ^63^Cu to CuCOX. *A*, Timeline of experiments for cells cultured under low Cu conditions and subsequently exposed to 30 µM ^63^Cu for 48 h (HA). Effects of the knockdowns of indicated transporters on the allocation of ^63^Cu to CuCOX (*B*), total Cu (*C*), Zn (D) and Fe (*E*). *F*, Timeline of experiments for cells cultured under 30 µM Cu conditions and subsequently exposed to 30 µM ^63^Cu for 48 h (HC). Effects of the knockdowns of indicated transporters on the allocation of ^63^Cu to CuCOX (G), total Cu (H), Zn (I) and Fe (*J*). K and L, western blots showing efficiency of the indicated knockdowns for HA (K) and HC (*L*) condition. M and N, RT-PCR validation of CTR1 knockdown under HA (*M*) and HC (*N*) conditions. Metals’ abundance was normalized to Phosphorus (P). Except for N and M, n=4 (biological replictaes); In M and N, n=3 (biological replictaes). Means±SD are shown; P values were calculated using two-tailed t-test and comparing significance of changes caused by specific siRNA to NT siRNA.

In the case of cells chronically grown continuously in low Cu and exposed to 30 µM ^63^Cu, the knockdown (KD) of CTR1, did not affect ^63^Cu accumulation and surprisingly somewhat increased the allocation of ^63^Cu to CuCOX in 786-O cells (Fig. 5*B*). In RCC4 cells, it had a small effect, decreasing ^63^Cu total content and did not affect the allocation of ^63^Cu to CuCOX (Fig. 5C). In contrast, there was a strong and consistent effect of DMT1 KD on ^63^Cu accumulation and allocation to CuCOX in both cell lines (Fig. 5, *B* and *C*). KD of LAT1 was effective in reducing total and CuCOX ^63^Cu in 786-O but not in RCC4 cells, supporting the role of two transporters in ^63^Cu uptake and CuCOX allocation, contributing to higher Cu concentrations in 786-O cells (Fig. 5, *B* and *C*). Finally, KD of SLC25A3 decreased allocation of ^63^Cu to CuCOX in both cell lines, but in 786-O cells had an additional effect on the total accumulation of ^63^Cu, potentially indicating a mechanistic link between allocation of Cu to CuCOX and an overall uptake of Cu (Fig. 5, *B* and *C*). Zn content was diminished by the KDs of all transporters in 786-O cells but only by DMT1 KD in RCC4 cells. Fe content was reduced by KDs of DMT1 and LAT1 in 786-O cells and was not affected in RCC4 cells. (Fig. 5, *D* and *E*). These effects were less strong in the case of Zn and Fe as compared to Cu.

The timeline of treatments with siRNAs and 30 µM ^63^Cu in the case of cells continuously grown in low Cu media was the same as shown in Fig. 5A (Fig. 5*F*). KDs of transporters in cells chronically exposed to high Cu and treated with 30µM ^63^Cu showed a significant contribution of DMT1 to the total Cu content and CuCOX, and Zn in 786-O cells and a contribution of LAT1 to the total levels of Cu but not to CuCOX (Fig. 5, *G* and *I*). There was no effect on the Fe concentrations (Fig. 5*J*). In the case of RCC4 cells, none of the KDs affected Cu, CuCOX, Zn or Fe (Figs. 5,*G-J*). The KDs of the transporters were validated by immunoblotting for DMT1, LAT1 and SLC25A3 (Fig. 5, *K* and *L*). Because there are no validated antibodies for CTR1, the KDs were demonstrated by RT-PCR (Fig. 5, *M* and *N*). Overall, these data support a major role of DMT1 in the uptake of Cu in renal cancer cells, while its role in the uptake of other divalent metals is more limited.

## Discussion

We established that the high molecular weight peak identified in SEC-ICP-MS chromatograms represents the active CuCOX complex and is regulated by Cu and metabolic conditions that support oxidative phosphorylation. This peak is induced within 48 h of Cu exposure, with extracellular Cu entering cells via transporters such as DMT1, LAT1, and mitochondria via SLC25A3. The induction of this peak is also associated with an increase in the oxygen consumption rate and mitochondrial ATP production, which are functional indicators of CuCOX activity. Copper uptake influences also the homeostasis of other essential metals, Zn and Fe. Notably, chronic exposure to elevated Cu levels is linked to increased intracellular Fe accumulation. This is especially relevant in the context of ccRCC, a cancer with high Cu content, which may also exhibit elevated Fe levels. Such a metal imbalance could sensitize cancer cells to ferroptosis, a regulated form of cell death driven by iron-dependent lipid peroxidation, when exposed to certain therapeutic interventions (28). This observation highlights a potential vulnerability that could be exploited in treatment strategies that target metal metabolism.

The measurement of CuCOX levels using SEC-ICP-MS in primary tumors or biopsies from both primary and metastatic lesions, combined with total Cu quantification in sera and tumor tissues by ICP-MS, could offer valuable clinical utility. This combined approach may serve as a prognostic biomarker for disease progression, providing insights into tumor oxidative metabolism and potentially guiding therapeutic decisions. The metallomic analysis of tumor Fe, another metal crucial for oxidative phosphorylation, is hindered by iron’s presence in a much larger number of proteins than Cu, particularly in high molecular mass fractions and its abundance in red blood cells.

Unexpectedly, our investigation revealed that exposing cancer cells to galactose not only induced CuCOX, as we initially hypothesized, but also led to a general increase in intracellular Cu levels. The mechanism by which galactose stimulates oxidative phosphorylation remains incompletely understood. While galactose is metabolized through the Leloir pathway into glucose (29), which then enters glycolysis to yield two ATP molecules, this process may occur at a slower rate compared to glucose metabolism. It has been proposed that this slower process reduces ATP availability, thereby promoting mitochondrial respiration as a compensatory mechanism. However, our findings nominate a novel pathway: galactose may stimulate mitochondrial respiration not through ATP limitation, but by enhancing Cu uptake and directing it toward CuCOX biogenesis, thereby boosting oxidative phosphorylation, OCR, and ATP production. This effect could be potentially mediated via galactosylation of membrane proteins or transporters.

Our results reveal novel insights into the role of copper transporters. Surprisingly, they suggest that the high-affinity Cu transporter CTR1 is not essential for Cu uptake under certain conditions and point to a robust mechanism of Cu uptake via DMT1 and LAT1. In particular, DMT1 appeared to be the most consistently involved in Cu import across experimental conditions, suggesting it may serve as a key mediator of Cu import. This finding not only enhances the current understanding of Cu trafficking in cancer cells but also implicates metal-ion transporters, traditionally associated with iron and amino acid metabolism, in regulating mitochondrial function through Cu delivery. Further investigation into the regulation and specificity of DMT1-mediated Cu uptake could reveal novel therapeutic targets, especially in Cu-dependent tumor types. Lastly, we observed differences in Cu uptake mechanisms when cells are chronically adapted to high Cu conditions. Here, DMT1 remains the primary transporter for Cu and Zn in 786-O cells, but its role in Fe uptake is limited, and the contributions of LAT1 and SLC25A3 are also diminished. In RCC4 cells, none of these transporters are necessary for Cu uptake, indicating that alternative or redundant mechanisms may exist. These differences likely stem from the need to limit Cu uptake and balance it with mechanisms that protect cells from Cu toxicity.

Our findings provide novel insights into the regulation of CuCOX activity in renal cancer cells, Cu transport mechanisms, and the potential vulnerabilities in metal metabolism that could be targeted in therapeutic strategies, particularly for Cu-dependent cancers like ccRCC.

### Experimental Procedures

### Cell culture and treatments

Human renal carcinoma cell lines 786-O (RRID: CVCL_1051, ATCC) and RCC4 (RRID: CVCL_0498) and human immortalized renal proximal tubule cells RPTEC cells TH1 (kerafast ECH001) (RRID: CVCL_K278) and TERT1 (ATCC, CRL-4031) were cultured in DMEM/F12 medium (Cytiva, SH30023) supplemented with 10% fetal bovine serum (FBS; Gibco, 6000-044) at 37°C in a humidified atmosphere containing 5% CO_2_. To generate high-copper (Cu) media, 300 μM CuSO_4_ was incubated with 100% FBS overnight at 4°C, followed by a 10-fold dilution into complete culture media. Chronic copper-high cells were generated by gradually increasing copper concentrations over a two-week period as described before (13). The 30 μM ^63^Cu-enriched media were prepared using Copper-63 metal (Cambridge Isotope Laboratories, CULM-463-PK). Media copper concentrations were routinely validated using ICP-MS. Cell line authentication was performed regularly by short tandem repeat (STR) profiling (LabCorp, Burlington, NC), and all cultures were screened for mycoplasma contamination using the MycoAlert™ Mycoplasma Detection Kit (Lonza). Cells were used for experiments up to passage 15.

### SEC-ICP-MS

Cell pellets were lysed in a buffer containing 1% digitonin, 0.1% SDS, 10 mM NaCl, and 50 mM Tris-HCl (pH 7.4), filtered through a 0.45 µm membrane, and injected into an Agilent 1290 HPLC system equipped with a thermostated autosampler (4°C), vacuum degasser, binary pump, column oven, and UV-Vis diode array detector. The size exclusion chromatography (SEC) column was a TSK gel QC-PAK GFC 200, 7.8 × 150 mm, 5 μm particle size (Tosoh Global) with 50 mM ammonium acetate and 0.5% MeOH at pH 7.4, at a flow rate of 0.565 mL min^-1^ and an injection volume of 80 µl. The outlet of the HPLC system was connected to the Agilent diode array UV-Vis detector with a 10 mm optical path to acquire full UV-Vis spectra from 220 to 750 nm every second, and then to the ICP-MS-MS nebulizer by a 65 cm PEEK capillary of 0.17 mm internal diameter. The ICP-MS was operated in time-resolved analysis under oxygen reaction mode (at 2 mL/min), with an integration time of 0.1 s per isotope, including ^63^Cu, ^65^Cu, ^66^Zn, ^57^Fe and ^31^P → ^47^SO isotopes. For quality control analysis, the column retention times were calibrated using a gel filtration standard (GFS, Bio-Rad Laboratories, 1511901). The intensity of the chromatographic peaks for Cu, Zn and Fe were normalized to the total phosphorus for each chromatogram. To analyze protein content at the individual chromatographic peaks, equal amounts of total cell lysates were injected for SEC-ICP-MS analysis, and fractions corresponding to CuCOX peaks were collected based on the retention time. Samples were then freeze-dried (Millrock, NY), resuspended in RIPA buffer and analyzed by immunoblotting.

The integration of the SEC-ICP-MS chromatograms was performed in the Origin X software package (OriginLab, MA) after exporting them from the Agilent Mass Hunter ICP-MS software. Quantification of the peaks was performed using Origin software and an in-house Excel script developed to compute the isotope dilution results. The exogenous ^63^Cu was determined using the ^65^Cu signal (in counts per second, CPS) at each point of the chromatogram to calculate the expected ^63^Cu signal based on the natural isotopic abundance ratio. This calculated ^63^Cu contribution was subtracted from the experimentally established ^63^Cu signal (in CPS). The remaining CPS values represent the excess of ^63^Cu at each point, enabling the generation of a new chromatogram specifically for exogenous ^63^Cu for analysis. Endogenous Cu was calculated as Total ^63^Cu minus Extra ^63^Cu. As a negative control, samples without ^63^Cu treatment were included resulting in a flat chromatogram. The areas under the chromatogram peaks were used to quantify the metal content in the SEC fractions. Quantification was performed by applying the sensitivity for ^63^Cu, obtained from a gel filtration standard with known Cu content. Finally, the metal content values were normalized to the total phosphorus in each sample.

The external calibration method was used to quantify the total Cu concentration of cell culture media and cell pellets as needed after an acid digestion in 10% nitric acid at 50ºC, in an Agilent 8900 ICP-MS/MS system (RRID: SCR_019460).

### Western Blots

Total cell lysates were prepared using RIPA buffer (Thermo Scientific, 89901) supplemented with 1× protease and phosphatase inhibitor cocktail (PPI; Thermo Fisher Scientific, 1861281) and 10 µM phenylmethanesulfonyl fluoride (PMSF; Sigma, P7626). Lysates were centrifuged at 14,000 × *g* for 20 min at 4°C. Protein concentration in the supernatant was quantified using the DC Protein Assay (Bio-Rad, 5000114). Equal amounts of protein were separated by SDS-PAGE.

For samples subjected to SEC-ICP-MS, fractions were collected from the HPLC system based on previously established retention times in SEC-ICP-MS. These samples were filtered, freeze-dried, and resuspended in RIPA buffer containing PPI and PMSF. Ten percent of the volume was used for Western blotting as described below.

Proteins were transferred to PVDF membranes, which were then blocked for 1 h at room temperature in 5% non-fat dry milk prepared in PBS-T (PBS + 0.1% Tween-20). Membranes were incubated with primary antibodies overnight at 4 °C with gentle rocking. Following three washes in PBS-T, membranes were incubated with HRP-conjugated secondary antibodies (1:5000 in 5% milk/PBS-T) for 1 h at room temperature. After additional washes, signals were developed using one of the following chemiluminescent substrates: Pierce ECL Western Blotting Substrate (Thermo Fisher Scientific, 32106), SuperSignal West Femto Maximum Sensitivity Substrate (Thermo Fisher Scientific, 34095), or SuperSignal West Atto Ultimate Sensitivity Substrate (Thermo Fisher Scientific, A38556). Blots were imaged using a Bio-Rad ChemiDoc system. Primary antibodies used: MT-CO1 (Abcam, ab203912; 1:1,000; RRID: AB_2801537); MT-CO2 (Abbexa, abx125706; 1:1,000; RRID: AB_3076223); COX4I1 (Cell Signaling Technology, 4844; 1:1,000; RRID: AB_2085427); GAPDH (Abcam, ab8245; 1:8,000; RRID: AB_2107448); DMT1 (Proteintech, 20507-1-AP; 1:1,000; RRID: AB_10694284); LAT1 (Cell Signaling Technology, 5347; 1:1,000; RRID: AB_10695104); SLC25A3 (Gift from P. Combine and S. Leary (34); 1:1,000). Secondary antibodies: Anti-Rabbit IgG HRP-linked (Cell Signaling Technology, 7073; 1:5,000; RRID: AB_2099233); Anti-Mouse IgG HRP-linked (Cell Signaling Technology, 7076; 1:5,000; RRID: AB_330924).

### RT-qPCR

Total RNA was extracted from cells using 1 mL of TriReagent (MRC, TR 118) following the manufacturer’s protocol. RNA was resuspended in 20 µL of nuclease-free water and quantified using a NanoDrop spectrophotometer; the 260/280 absorbance ratio was recorded to assess purity. Complementary DNA (cDNA) was synthesized using the High-Capacity cDNA Reverse Transcription Kit (Applied Biosystems, 4368814). Quantitative PCR was performed using the ΔΔCt method with 1× Fast SYBR Green Master Mix (Applied Biosystems) on an Applied Biosystems QuantStudio 7 Real-Time PCR System (RRID: SCR_020245), using 20 ng of cDNA per reaction. Gene expression was normalized to the housekeeping gene PP1A, and expression levels were calculated relative to the Non-Targeting control sample. Primer sequences were as follows: CTR1 (SLC31A1): Forward: 5′-CCC TTA CTC TGT TGT CCT TTC-3′ Reverse: 5′-CAC AGC ATA GCA CTG TCT AC-3′; PP1A: Forward: 5′-ACC GCC GAG GAA AAC CGT GTA-3′ Reverse: 5′-TGC TGT CTT TGG GAC CTT GTC TGC-3′.

### siRNA Transfections

Cells were transfected with small interfering RNA (siRNA) using Lipofectamine 3000 Transfection Kit (Invitrogen, L3000015) following the manufacturer’s instructions. siRNAs were used at a final concentration of 50 nM. Control cells were treated with non-targeting siRNA. Culture media were replaced 24 h after the first transfection. A second siRNA transfection (50 nM) was performed the following day. Twenty-four hours after the second transfection, cells were treated with ^63^Cu-containing media for 48 h. The following siRNA pools were used: Non-Targeting pool (Dharmacon, D-001810-10-20); SLC31A1 (CTR1): Dharmacon, L-007531-02-0005; SLC11A2 (DMT1): Dharmacon, L-007381-00-0010; SLC7A5 (LAT1): Dharmacon, L-004953-01-0005; SLC25A3: Dharmacon, L-007484-00-0010.

### Seahorse Metabolic Assays

Oxygen consumption rate (OCR) and extracellular acidification rate (ECAR) were measured using an Agilent Seahorse XFe96 Analyzer. For galactose treatment conditions, cells were pre-treated for 48 hours in DMEM (Gibco, A14430-01) supplemented with 10% dialyzed fetal bovine serum (dFBS), 10 mM glucose or 10 mM galactose, and 4 mM glutamine. On the day of the experiment, cells were incubated for 1 h in Seahorse XF Base Medium (Agilent) without serum, containing 10 mM glucose or 10 mM galactose and 4 mM glutamine. Basal respiration, maximal respiration, and ATP-linked OCR were determined using the Agilent Seahorse Mitochondrial Stress Test. Measurements were taken before treatment and after sequential injections of: Oligomycin A (1.5 µM); FCCP at cell line-specific concentrations: 786-O: 0.25 µM, RCC4: 1 µM, TH1: 1 µM, TERT1: 0.5 µM, Antimycin A/rotenone (0.5 µM each). Data were analyzed using the Agilent Mitochondrial Stress Test Report Generator and normalized to cell number using Hoechst fluorescence staining. To determine mitochondrial vs. glycolytic ATP production, OCR and ECAR were measured before treatment and after sequential injections of oligomycin A (1.5 µM), followed by antimycin A/rotenone (0.5 µM each). Data were analyzed using the Agilent ATP Rate Assay Report Generator.

## Statistics

Significance was determined using two-tailed unpaired t-test. Box and whisker plots displayed the median, minimum and maximum values, as well as all data points. Bar graphs displayed the mean with standard deviations, and all data values.

## Data availability

All metalomics data can be provided upon request from the corresponding authors.

## Acknowledgments

The authors thank Addison Cooper (Cooper Graphics) for preparing the figures.

## Funding and additional information

The following grants supported the work: MCK (R01CA287260 and 2I01BX001110 BLR&D VA Merit Award); KEV (R35GM146878); MEB (T32CA17846); DS (T32ES007250). The content is solely the responsibility of the authors and does not necessarily represent the official views of the National Institutes of Health.

